# Thermodynamic analysis of the GAS_right_ transmembrane motif supports energetic model of dimerization

**DOI:** 10.1101/2022.07.11.499632

**Authors:** Gladys Díaz Vázquez, Qiang Cui, Alessandro Senes

**Affiliations:** Department of Biochemistry, University of Wisconsin-Madison, Madison, WI 53706; Biophysics Graduate Program, University of Wisconsin-Madison, Madison, WI 53706; Department of Chemistry, Boston University

## Abstract

The GAS_right_ motif, best known as the fold of the glycophorin A transmembrane dimer, is one of the most common dimerization motifs in membrane proteins, characterized by its hallmark GxxxG-like sequence motifs (GxxxG, AxxxG, GxxxS, and similar). Structurally, GAS_right_ displays a right-handed crossing angle and short inter-helical distance. Contact between the helical backbones favors the formation of networks of weak hydrogen bonds between Cα–H carbon donors and carbonyl acceptors on opposing helices (Cα–H∙∙∙O=C). To understand the factors that modulate the stability of GAS_right_, we previously presented a computational and experimental structure-based analysis of 26 predicted dimers. We found that the contributions of van der Waals packing and Cα–H hydrogen bonding to stability, as inferred from the structural models, correlated well with relative dimerization propensities estimated experimentally with the *in vivo* assay TOXCAT. Here we test this model with a quantitative thermodynamic analysis. We used FRET to determine the free energy of dimerization of a representative subset of 7 of the 26 original TOXCAT dimers using FRET. To overcome the technical issue arising from limited sampling of the dimerization isotherm, we introduced a globally fitting strategy across a set of constructs comprising a wide range of stabilities. This strategy yielded precise thermodynamic data that show strikingly good agreement between the original propensities and ΔG° of association in detergent, suggesting that TOXCAT is a thermodynamically driven process. From the correlation between TOXCAT and thermodynamic stability, the predicted free energy for all the 26 GAS_right_ dimers was calculated. These energies correlate with the *in silico* ΔE scores of dimerization that were computed on basis of their predicted structure. These findings corroborate our original model with quantitative thermodynamic evidence, strengthening the hypothesis that van der Waals and Cα–H hydrogen bond interactions are the key modulators of GAS_right_ stability.

**Secondary Abstract:** We present a thermodynamic analysis of the dimerization of the GAS_right_ motif, a common dimerization motif in membrane proteins. Previously, we found that the stability of GAS_right_ is modulated by van der Waals packing and weak hydrogen bonds between Cα–H carbon donors and carbonyl acceptors on opposing helices. The experimental dimerization propensities were obtained with an *in vivo* assay. Here we assess this model quantitatively by measuring the free energy of dimerization of a subset of the original dimers. The thermodynamic data show strikingly good agreement between the original propensities and their ΔG° of association, confirming the model and strengthening the hypothesis that van der Waals and Cα–H hydrogen bond interactions are the key modulators of GAS_right_ stability.

## Introduction

Membrane protein oligomerization is a fundamental process in the life of a cell. Oligomerization is especially important for bitopic proteins, i.e. the membrane proteins that contain a single transmembrane (TM) helix. The association of these TM helices can be optimized for stability in constitutive dimers, such as the case of glycophorin A (GpA)^1^. In other instances, stability is tuned appropriately to support dynamic association, which can be critical for regulating signal transduction or activation in important biological systems, such as integrins^2,3^ and receptor tyrosine kinases^4–6^, to name a few. Understanding the interplay between the forces involved in TM helix oligomerization could support the prediction of structure and stability, the identification of potential conformational changes, and the interpretation of the effect of mutations in these system, providing insight into biological function and regulation.

In the present study, we focus on the energetics of association of an important structural dimerization motif known as GAS_right_^7^, which is best known as the fold of the prototypical GpA dimer^1^. The GAS_right_ motif is one of the structural motifs most commonly observed in dimeric transmembrane complexes^8,9^. It is characterized by the presence of the small amino acids, <u>G</u>lycine, <u>A</u>lanine, and <u>S</u>erine (GAS) at the dimer interface, and a <u>right</u>-handed crossing angle of approximately −40° between the two helices. The small residues are separated by three amino acids and arranged on the same face of the helix to form GxxxG-like sequence motifs (GxxxG, GxxxA, SxxxG, etc.)^10–12^ (Fig. 1). They form a flat face that allows the helical backbones to come in close contact, promoting tight packing. The contact between the backbones at the specific geometry of GAS_right_ promotes the formation of networks of weak hydrogen bonds between Cα–H carbon donors and carbonyl acceptors on opposite helices (Cα–H∙∙∙O=C)^7^. C–H groups are typically weak hydrogen bond donors unless they are activated. In the case of the Cα–H group, the flanking amide groups act as electron-withdrawing substitutes with respect to the Cα carbon, which, in turn, results in significant polarization of the Cα–H bond. Indeed, quantum mechanics calculations have estimated the energy of Cα–H hydrogen bonds in proteins to be approximately one-half of that of N–H donors in vacuum^13,14^. For this reason, they are likely to contribute significantly to the free energy of dimerization of GAS_right_ dimers, considering that multiple instances of these hydrogen bonds occur at the same interface in this motif^7,15^.

**Fig. 1.**
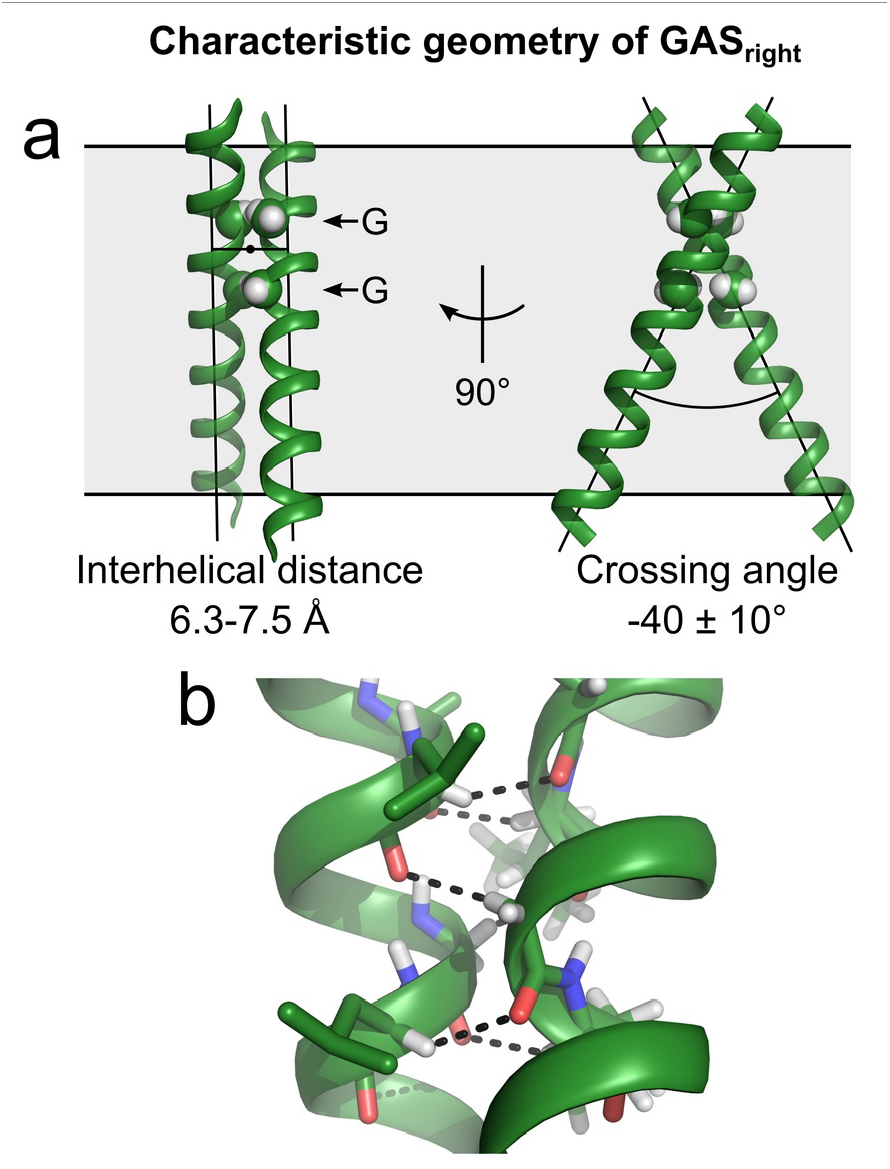
Geometry of the GAS_right_ motif. *a*) GAS_right_ is a right-handed helical dimer with a short inter-helical distance (6.3-7.5 Å) and a right-handed crossing angle of approximately −40°. The GxxxG sequence pattern at the crossing point (indicated by the arrows) allows the backbones to come into contact. *b*) This contact enables formation of networks of 4 to 8 weak inter-helical H-bonds between Cα–H donors and carbonyl oxygen acceptors at the crossing point of the dimer.

To test the hypothesis that Cα–H hydrogen bonds, along with van der Waals (VDW) packing, are major drivers of stability in GAS_right_ dimers, we previously used a combined computational experimental approach^16^. Specifically, we predicted the structure of a series of 26 GAS_right_ dimers using the program CATM^7^ and compared the energy score that was calculated with the experimental dimerization propensities obtained with TOXCAT^17^, a genetic assay that measures oligomerization in the *Escherichia coli* membrane (Fig. 2*a*). To reduce some of the variability that is typical of a biological assay, we redesigned the constructs by “stitching” the 8 positions predicted by CATM to be at the dimer interface into a standardized 21 amino acids poly-Leu backbone (LLLxxLLxxLLxxLLxxLILI, where the x represents the variable interfacial amino acids)^16^. Such standardization was important to reduce the differences in protein expression level. It also allowed us to focus on the forces that play a role at the interaction interface, isolating them from other variables that could contribute to dimerization stability, such as the length of the hydrophobic region of the TM helices and the position of the crossing point in the dimer, which were features shared by all constructs (Fig. 2*b*). The predicted structures were partially validated by mutating the critical interfacial glycine at C1 (the position at the interface that is in the closest contact with the opposing backbone) to a large isoleucine, a mutation that is expected to completely abolish dimerization by creating steric clashes^16^.

**Fig. 2.**
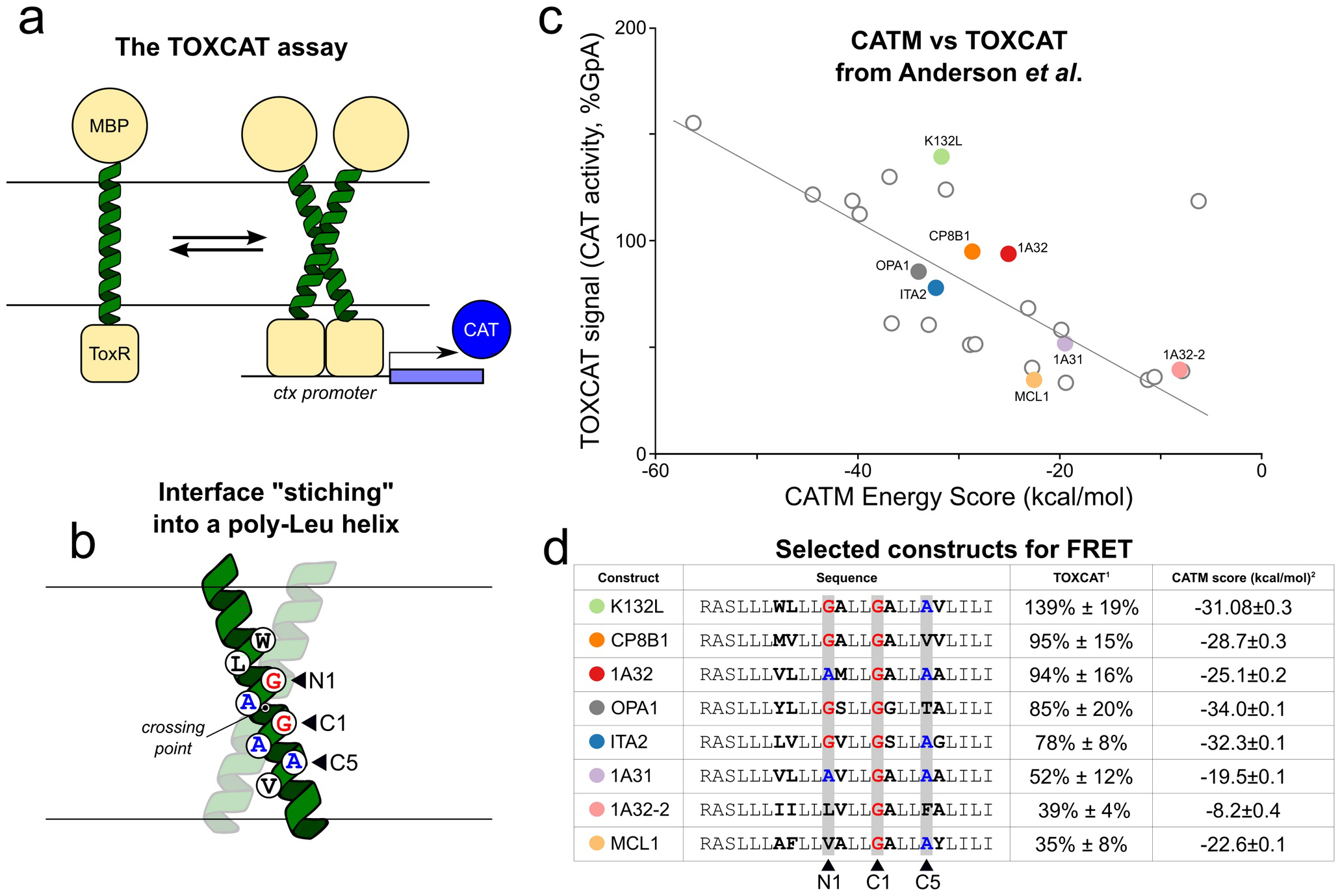
Construct selection. a) In our previous study^16^, we used TOXCAT to study the dimerization of a set of predicted GAS_right_ dimers. TOXCAT is an *in vivo* reporter assay that provides propensities of dimerization relative to a known standard. In the assay, the TM domain under investigation is fused to the ToxR transcriptional activator. TM helix association results in the expression of a reporter gene in *E. coli* cells (CAT), which can be quantified. b) To reduce variability in TOXCAT, the eight amino acids that formed the interface as predicted by CATM were “stitched” into a standardized poly-Leu sequence. c) In our previous study, we found a correlation between the energy score predicted with the program CATM and the dimerization propensity measured with TOXCAT assay for a series of 26 standardized GAS_right_ dimers. Plot reproduced here from Anderson *et al*.^16^. The data suggest that two of the primary interactions that contribute to modulating the dimerization stability of these constructs are a combination of VDW interactions and weak Cα–H hydrogen bonding. The 8 constructs selected for the present study are highlighted in color. d) List of the subset of eight GAS_right_ dimers analyzed *in vitro* in this study to measure their ΔG° of dimerization. The subset covers a range of TOXCAT homodimerization propensities and CATM energy scores.

As shown in Fig. 2*c*, in our previous study we identified a statistically significant relationship between experimental stability estimated with TOXCAT and the computational energy scores of the constructs (data reproduced from Anderson *et al*.^16^). However, neither CATM nor TOXCAT are rigorous methods for assessing the free energy of association of TM dimers. CATM is a program that can predict the structures of known GAS_right_ dimers with high accuracy, but its energy function is relatively simple, consisting of an unweighted sum of only three terms: VDW, hydrogen bonding, and an implicit solvation function. The ΔE score it produces is obtained by calculating the differential of these three terms between static monomeric and dimeric models. TOXCAT reports relative propensities of association relative to know standards (glycophorin A and its monomeric G83I variant), but it does it indirectly, through the stimulation of the expression of a reporter gene (Fig. 2*a*). It indicates semi-quantitatively whether a dimer is strong, moderately stable or weak but it does not provide free energies of association. The assumption is that the signal is proportional to the amount of dimer present in the membrane, and thus the response is thermodynamically driven ^18^, an assumption that was yet to be fully tested. Furthermore, the assay could be affected by the variability of a living biological system and in particular by differential expression levels of the constructs. Despite these limitations, by sampling a large set of validated GAS_right_ constructs of different stabilities with these two methods, in our previous study we were able to identify features that correlated statistically with the apparent stability of the dimers^16^. The data indicated that packing and Cα–H hydrogen bonds were major factors that modulated the stability of these constructs and suggested that stability can be governed by inter-helical geometry, which in turn is enabled by the sequence^16^.

In this study, we take an important step forward towards formulating a quantitative model of the energetics of GAS_right_, with a quantitative thermodynamic analysis of dimerization *in vitro*. We selected a representative subset of the original 26 GAS_right_ dimers^16^ and used Fӧrster Resonance Energy Transfer (FRET) to measure their free energy of association in detergent. We found a strong correlation exists between the free energy of association in detergent and the dimerization propensity obtained previously with TOXCAT. The data also suggest that there is a striking correspondence between the ΔΔG° of association measured *in vitro* and the same quantities in the membrane of *E. coli*. The study provides a thermodynamic foundation for the model that identifies a combination of VDW packing and weak hydrogen bonding as the primary drivers and modulators of stability of GAS_right_ dimers.

## Results and Discussion

### FRET construct design

While the majority of oligomerization studies of TM helices based on FRET have been performed on synthetic peptides^19–23^, we opted for biological expression to increase throughput. Specifically, we used a chimeric construct that fuses a soluble Staphylococcal nuclease (SN) domain to the TM helices, a strategy used in previous thermodynamic association studies based on other methods^24–31^. As a minor change, we added a 15-amino acid long flexible linker between the TM helix and SN moieties, to structurally decouple the orientation of the two domains and thus minimize any influence of the geometry of the TM region on the FRET efficiency. For labeling, we screened three surface positions, including an isosteric mutation (S3C), and two Ala positions that were previously found not to dramatically affect SN’s stability when changed to cysteine (A60C and A112C)^32^. Variant A112C yielded calculated labeling efficiencies of 70-80% (supplementary Table S1) when reacted with either Cy5-maleimide (acceptor), or Cy3-maleimide (donor) and was thus selected for the study (Fig. 3a).

**Fig. 3.**
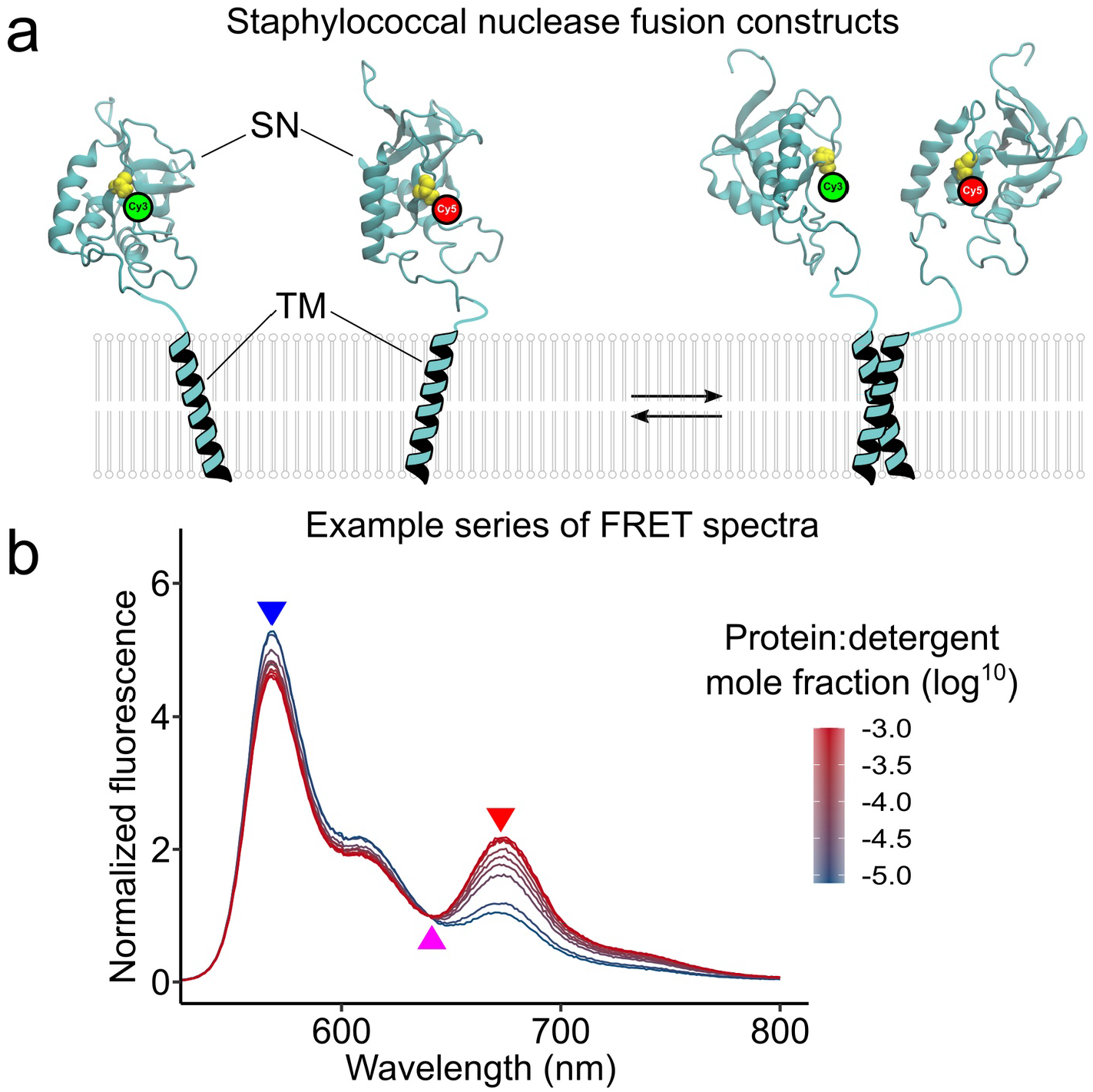
The FRET experiment. a) The reaction under study corresponds to a monomer-dimer equilibrium. The N-terminus of the TM helices of interest were fused to the C-terminus of the soluble staphylococcal nuclease (SN) through a 15-residues flexible linker. SN was labeled at Cys residue introduced at position 112 with either Cy3-maleimide (donor) or Cy5-maleimide (acceptor) for FRET experiments. b) Example of a series of fluorescence spectra obtained at different mole fractions (moles of protein/moles of detergent), driv ing the reaction from a monomeric to a dimeric state (construct SN-GpA). The fluorescence spectra were normalized using the isosbestic point indicated by the magenta arrow at 640 nm. The donor emission at 570 nm (blue arrow) decreases and the acceptor emission at 670 nm (red arrow) increases as the protein mole fraction is increased. Original and normalized spectra of all constructs are shown in supplementary Fig. S1.

From the 26 constructs from our previous TOXCAT study^16^ we excluded five that contained a cysteine in their sequence to avoid possible issues with non-specific labeling. The constructs that did not clone efficiently or expressed poorly were also eliminated. This yielded a final set of eight constructs that cover a wide range of homodimerization propensities in TOXCAT and energy scores predicted by CATM (Fig. 2d). These constructs were expressed, purified in *n*-decyl-β-D-maltopyranoside (DM) detergent, and labeled.

### Theoretical framework of the FRET experiments and global fitting analysis

FRET experiments were performed by keeping the amount of protein constant and changing the concentration of the DM detergent in solution, under the well-established assumption that the equilibrium of the dimerization reaction is governed not by the total volume but by the “hydrophobic volume”, given that the TM helices are constrained to reside in the interior of a detergent micelle^33^. Protein concentration is thus expressed as protein:detergent mole fraction, which was explored over a range of approximately 1:10 ^−5^ to 1:10^−3^ to drive the reaction from monomer to dimer states. The mole fraction was corrected by subtracting the critical micellar concentration from the detergent concentration, to account for the presence of monomeric (non-micellar) detergent.

An example series of donor-acceptor fluorescence spectra at increasing protein mole fraction is illustrated in Fig. 3*b*. The spectra were first corrected by subtracting the contribution to emission of acceptor-only samples with the same concentration and then normalized using the isosbestic point (640 nm for our system), to correct for changes in signal that are not related to the transfer of energy between the fluorophores, such as small errors in the estimate of concentration as well as differences in labeling efficiencies^34^. Original and normalized spectra of all constructs are shown in supplementary Fig. S1.

The FRET series was validated by plotting donor emission at 570 nm against acceptor emission at 670 nm (supplementary Fig. S2). For all constructs that cover the transition from mostly monomeric to mostly oligomeric (GpA, ITA2, K132L, 1A32, CP8B1, and OPA1) we observed an inverse linear relationship, with consistent slopes and intercept values across the constructs, as expected when changes in fluorescence intensity are primarily due to FRET. Lower R^2^ are to be unexpected for 1A32-2 and other monomeric or weakly associating variants (G83I,1A31, and MCL1) because their data points are clustered in the upper left corner, in the low-FRET region, and do not cover as much of a spread for linear regression.

Equilibrium constants of dissociation were obtained by fitting the normalized fluorescence intensity at 670 nm (*NF*_670_) as a function of the protein:detergent mole fraction (*χ_T_*) using equation 1 (derived in supplementary material):

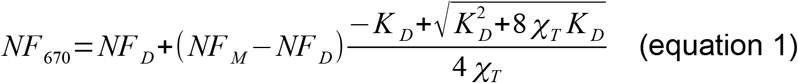

Three parameters were derived from the fitting. Two of them were global parameters, *NF_D_* and *NF_M_*, i.e. the normalized fluorescence intensities at the limits in which the samples are fully dimeric (at infinite concentration) or monomeric (at infinite dilution), respectively. The third parameter was the desired dissociation equilibrium constant *K_D_*, which was individually fitted to the data of each construct.

A major challenge presented by this type of FRET analysis is obtaining enough coverage of the association range (from mainly monomeric to mainly dimeric) so that all three parameters can be calculated with sufficient confidence. As illustrated in Fig. 4, the weakest dimers (MCL1, 1A32-2, and 1A31) cover primarily the monomeric region, whereas the strongest dimers (CP8B1 and the control GpA) cover most of the transition and approach a fully dimeric state but lack the monomeric baseline. Fitting the *NF_D_* and *NF_M_* parameters globally solves this problem since these baselines are expected to be similar across all constructs.

**Fig. 4.**
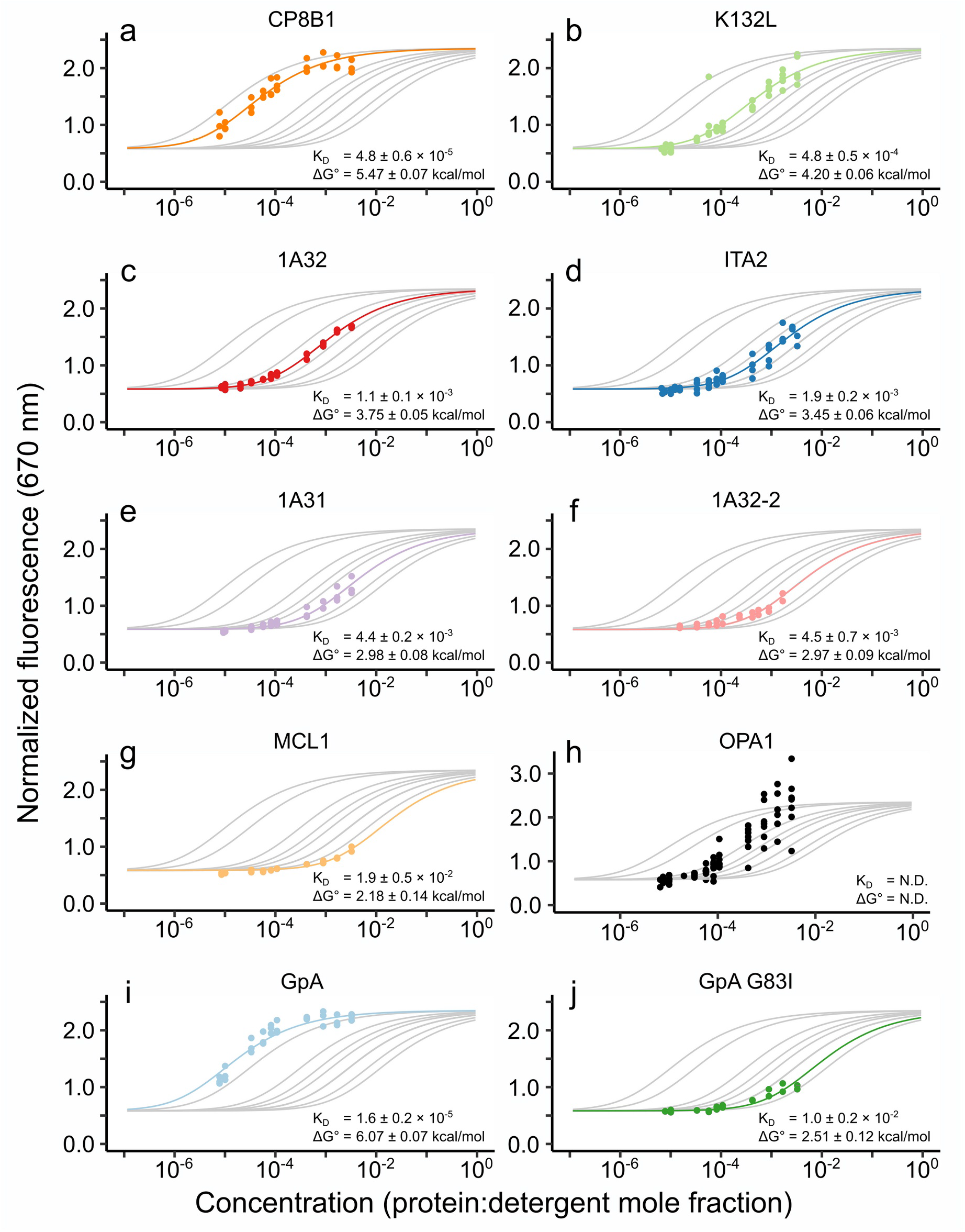
Thermodynamic analysis, global fitting of FRET data. The graphs show the fluorescence data as a function of concentration in FRET experiments for the eight selected GAS_right_ constructs from the 26 previously studied using TOXCAT (*a*-*h*). The prototypical GAS_right_ dimer GpA (*i*) and its monomeric mutant GpA G83I (*j*) were also included as controls. The gray lines correspond to the global fit of the FRET data, which yielded the dissociation constants (K_D_). The construct OPA1 (*h*) was excluded from the global fitting because it showed a larger emission range and its stoichiometric analysis indicated it forms higher-order oligomers (supplementary Fig. S3).

It should be noted that the treatment requires the labeling efficiencies of the donor to be consistent across samples, as well as those of the acceptor (however, the labeling efficiencies of the donor and acceptor do not need to be identical). The normalization by the isosbestic point takes care of differences if labeling differences occurs in a limited range, particularly of the labeling efficiency of the donor and the establishment of the monomeric baseline *NF_M_*. In theory, a second rescaling term should be introduced to correct for the relative variation in acceptor efficiency relative to the donor (the Cy5% / Cy3% ratio), which affect more the dimeric baseline *NF_D_*. However, we found that this second factor did not improve our global fitting results, and it thus was not possible to determine whether it is beneficial when these ratios are sufficiently close.

In practice, inconsistent labeling can often be mitigated simply by dilution. Highly labeled samples can be diluted with unlabeled protein, bringing the set of samples closer to a common value. It is also possible to combine donor and acceptor samples of the same construct in different ratios than 50:50, to increase the labeling of one species and decrease the other. For example, if the donor’s efficiency is 90% and the acceptor’s efficiency is 60%, mixing them in a 4:6 ratio would bring them both at a 72% labeling level. It should also be kept in mind that for good quality FRET data, ideally, all samples should be preferably >70% labeled for both donor and acceptor.

### The selected GAS_right_ dimers cover a wide range of thermodynamic stabilities in vitro

Fig. 4*a*-*h* shows the normalized fluorescence data as a function of mole fraction for the eight selected GAS_right_ constructs. It also shows the fitted binding curves and the calculated free energies of dissociation (with the exception of construct OPA1, panel *h*, which was not included in the fitting). Two additional controls were also measured: the highly stable GpA dimer (panel *i*) and its monomeric G83I variant (*j*). Inclusion of these two controls of known stability contributed to establishing the *NF_D_* (wild type GpA) and *NF_M_* (G83I variant) baselines in the global fitting analysis.

Overall, the constructs cover a wide range of thermodynamic stabilities with ΔG° values ranging approximately from 2 to 6 kcal/mol. There is good consistency among the baselines of the various samples. Specifically, the constructs that occur in a prevalent unassociated state at the highest dilutions (ITA2, K132L, G83I, 1A32-2, 1A32, MCL1, and 1A31) are all in good agreement with the globally fit monomeric baseline (*NF_M_* = 0.58 ± 0.01). Correspondingly, the two constructs that approach a fully associated state at their highest concentrations (GpA and CP8B1) are consistent with the calculated dimeric baseline (*NF_D_* = 2.35 ± 0.03). Finally, the data points of the construct that span a significant portion of the monomer/dimer transition (GpA, CP8B1, ITA2, K132L, and 1A32) fit the curves well. Only the construct OPA1 displayed a larger emission range than the others, suggesting it may form higher-order oligomeric states. This was confirmed by stoichiometry analysis by varying donor-acceptor ratio^35,36^ (supplementary Fig. S3). For this reason, OPA1 was not included in the global fitting and was no longer considered in the analysis.

The standard association free energy that we obtained for GpA was −6.07 ± 0.07 kcal/mol. This value is comparable with previous thermodynamic studies of GpA in other detergents by analytical ultracentrifugation, which reported free energies of −5.7 ± 0.3 and −9.0 ± 0.1 kcal/mol in C14 betaine^27^ and C_8_E_5_^26^, respectively. As expected, the GpA G83I monomeric variant is much destabilized (−2.51 ± 0.12 kcal/mol). These results demonstrate that the global fit strategy across a set of constructs of varied stability enables the determination of the dissociation constant for a series of samples that individually do not cover a sufficient range from dissociated to associated states.

### Strong correspondence of relative stabilities in detergent and the *E. coli* membrane

After obtaining the thermodynamic stability of the constructs in detergent *in vitro*, we compared it with their dimerization propensities in biological membranes obtained previously with TOXCAT^16^. As shown in Fig. 5*a*, we observed the expected non-linear monotonic inverse relationship between the ΔG° of association and TOXCAT signal (Spearman’s rank correlation, ρ = 0.96, *p*-value = 0.003). Only one notable outlier does not follow the proportionality, CP8B1, which is the most stable among the constructs in this study but displayed only intermediate stability in the TOXCAT study. Among all constructs of our previous TOXCAT study, CP8B1 was the one that displayed the largest variation in its degree of expression (Fig S2 in Anderson et al.^16^), therefore this discrepancy may be possibly due to significant differences in protein concentration of CP8B1 in the *E. coli* membrane compared to the other constructs.

**Fig. 5.**
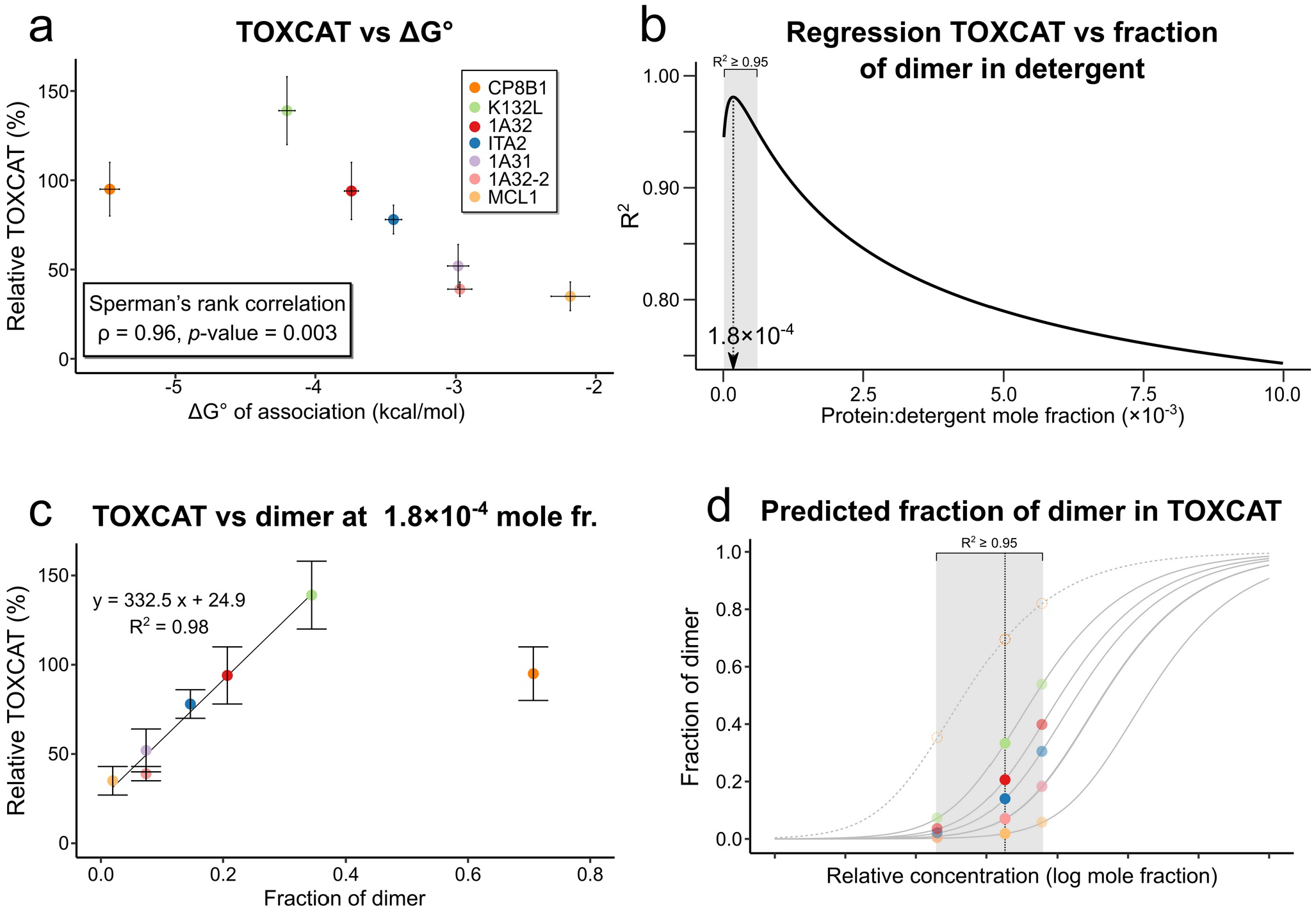
Comparison of the FRET results with the previous TOXCAT analysis. a) The thermodynamic stabilities of the constructs obtained here in detergent are compared to their previously determined TOXCAT homodimerization propensities. The expected inverse non-linear relationship is observed. A clear outlier in the trend can be observed, corresponding to CP8B1. This outlier was included in the statistical analysis. b) Results of the linear regression analysis between TOXCAT and fraction of dimer calculated at different mole fractions. Here the regression coefficient R^2^ is plotted (linear fits plotted in supplementary Fig. S4). The coefficient is maximized at a 1.8×10^−4^ mole fraction. The region in which R^2^ ≥ 0.95 is shaded, corresponding to concentrations between 2×10^−5^ and 6×10^−4^ mole fractions. The outlier CP8B1 was not included in this linear fit. c) Linear fit of TOXCAT vs fraction of dimer at 1.8 × 10^−4^ mole. The outlier CP8B1 was not included in this linear fit. d) The predicted fraction of dimer in the biological membrane of *E. coli*, based on the linear relationship observed between TOXCAT and the fraction of dimer, under the assumption that CAT expression is directly proportional to fraction dimer. The data suggest the constructs are mostly monomeric in TOXCAT, ranging from 2% for the weakest dimer (MCL1) to 35% for the strongest (K132L) from the conditions of maximum correspondence with the free energy data. The region that covers the predicted fraction dimer for all concentrations that produced an R^2^ ≥ 0.95 is also indicated (shaded).

The overall good correlation with the FRET data suggests that the response of TOXCAT is indeed governed by the thermodynamic stability of the dimers and that the expression of the CAT reporter gene is directly proportional to the amount of dimer that occurs in the membrane^16^. It also suggests that there is a close relationship between the stability of the constructs in detergent and the membrane. This relationship cannot be resolved directly from the TOXCAT data because of two unknown quantities. First, the free energies of dimerization of the constructs in the two environments are likely different^37^. In addition, the concentration of constructs in the *E. coli* membrane is unknown. However, if we make the assumption that the expression level in TOXCAT is relatively similar across this set of standardized poly-Leu constructs (as assessed by Western blotting^16^), we can ask whether the relative stability (ΔΔG°) of the series of constructs is conserved between the two environments.

We asked this question by checking if there is a correlation between the amount of dimer in detergent and their TOXCAT signal, which we assume is a proxy for the amount of dimer in the membrane. To do so, we performed a series of linear regressions between TOXCAT signal and the dimer fraction in detergent calculated across a range of mole fractions that cover the entire transition from fully monomeric to fully dimeric, as shown in supplementary Fig. S4 (the construct CP8B1 was excluded from this analysis since it is a clear outlier in the TOXCAT vs stability relationship).

We found that TOXCAT and dimer fraction correlate extremely well at certain concentration regimes. The regression coefficient plotted against protein:detergent mole fraction reaches a maximum of R^2^ = 0.98 at a mole fraction of 1.8×10^−4^ (Fig. 5*b,c*). At this mole fraction, the amount of dimer ranges from approximately 2% for the weakest dimer to 35% for the strongest (Fig. 5*d*). It should be noted that the correlation remains very strong for a range of concentrations. As a reference, the range in which R ^2^ ≥ 0.95 is between 2×10^−5^ and 6×10^−4^ mole fraction (shaded region in Fig. 5*b*). We conclude that the monomer/dimer equilibria of the six constructs occur in the membrane at levels of dimerization that are close to the regime observed at 1.8×10^−4^ in detergent (dashed line in Fig. 5*d*) and most likely falling on the left half of the binding curve (the shaded area in Fig. 5*d*). In this region, most constructs go from all nearly monomeric to reaching halfway in the saturation curve for the most stable. These data, therefore, are in reasonable agreement with a previous suggestion that TOXCAT constructs exist in mostly monomeric state in the membrane^18^.

These results indicate that there is a striking correspondence between relative stability (ΔΔG°) in detergent and the *E. coli* membrane for this set of constructs. They confirm for the hypothesis that the TOXCAT process is governed by thermodynamic stability^18^. Finally, the findings provide validation for the experimental data that was used to base the model that a combination of VDW packing and weak hydrogen bonding act as the primary drivers and modulators of stability of GAS_right_ dimers^16^.

### The CATM energy score captures predicted stabilities in detergent

After examining the relationship with TOXCAT, we proceeded to compare the free energy of association *in vitro* with the energy score we previously calculated with the structural prediction program CATM^16^. We first compared the experimental ΔG° of association in detergent of the subset of seven GAS_right_ constructs analyzed here with their CATM energy score (Fig. 6*a*). The linear regression did not produce a significant correlation (R^2^=0.26 *p*-value=0.2446), indicating that CATM does not capture well the energetics of association of this particular subset of constructs. We then took advantage of the derived relationship between TOXCAT signal and the ΔG° of association to back-calculate predicted ΔG° values in detergent for all 26 original constructs (supplementary Table S2) and assess the correlation with their CATM energy score. In this case, we found a reasonable correlation (R^2^=0.46) with highly significant *p*-value = 0.0001354 (Fig. 6*b*). The majority of the points form a clear trend dispersed around the line, with only two data points performing as clear outliers (TNR12, ROMO1). This discrepancy between Fig. 6*a* and *b* is not unreasonable: although a clear correlation can be identified with a larger set of 26 constructs, the same correlation may not become apparent with a smaller number of data points due to the noisy relationship and the loss of statistical power.

**Fig. 6.**
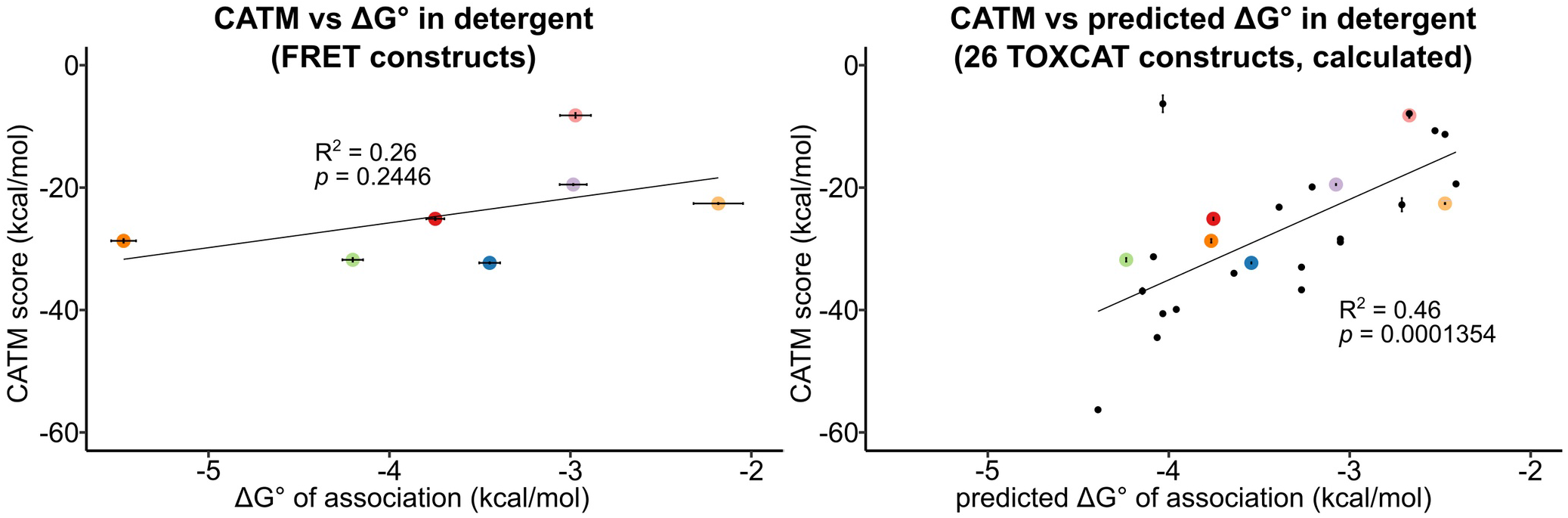
Comparison of experimental and computational energies of dimerization. a) The experimental ΔG° of association for the 7 constructs studied here do not result in a significant linear correlation with the CATM energy scores. b) The comparison was expanded by predicting the free energy of association for all 26 original constructs from their TOXCAT data. A significant correlation between this set and the CATM energy score was observed.

It should be noted that in Fig. 6*b* there is a large difference in the scale of the two axes, with approximately 17 kcal/mol of CATM energy corresponding to just 2 kcal/mol of predicted ΔG° in DM. This difference is not straightforward to interpret physically. It appears that CATM somehow over-estimates the interactions, highlighting a need for recalibrating its energy function, which is not unexpected. The CATM function is simple, as an unweighted sum of only three terms, VDW interactions (CHARMM 22^38^), hydrogen bonding (SCRWL 4^39^), and implicit solvation (IMM1^40^). The score ignores other potentially important terms, such as, for example, electrostatics or entropic contributions. With the availability of a sufficiently large data set of validated GAS_right_ TOXCAT constructs, it would be possible to derive (and rigorously test) an effective energy function to predict the relative stability of these dimers in detergent and potentially in the biological membrane. Nevertheless, it is notable that the simple energy function of CATM in its current form, appears to be a reasonable predictor of the relative stability of a large set of GAS_right_ dimers, confirming the model.

## Conclusions

Understanding the physical basis of folding and association in membrane proteins remains challenging because of the technical difficulties posed by these systems, together with the complexity of a process occurring in an anisotropic and highly heterogeneous milieu^41^. Many factors have been proposed to play a role in membrane protein folding and association. Among them are lipid-specific effects (such as solvophobic exclusion, specific lipid binding, lateral pressure, and hydrophobic matching^42–49^) and a variety of physical interactions (for example, aromatic and cation-π interactions^50–52^). However, two prominent candidates have emerged as the major determinants of stability in membrane proteins, namely the packing of apolar side chain and hydrogen bonding.

Packing is ubiquitous at helix-helix interfaces. These helices consist primarily of hydrophobic amino acids and their packing results in extensive and favorable VDW interactions^41,44,53,54^. Structural informatics indicate that side chains pack more efficiently in membrane proteins^54–57^ whereas mutations that disrupt packing are generally strongly detrimental to stability^18,53,58,59^, which clearly indicates that good packing is a necessary feature of stable membrane proteins and complexes. The question is to what extent packing can be a main driving force beyond a mere necessity. The favorable packing of apolar amino acids at helical interfaces in the associated state is counterbalanced by the favorable VDW interactions that the amino acids form with the lipids acyl chains in the unassociated state. Therefore, for packing to be an actual driving force, the protein-protein interactions in the folded state need to overcome the contributions of protein-lipid interactions that are lost upon folding. Although this question remains open, the recent design of a stable oligomeric complex created solely around apolar packing offers proof-of-principle evidence that is consistent with VDW interactions as an actual driving force^60^.

The second main candidate that has emerged as an important force for membrane protein folding is hydrogen bonging. There is little doubt that “canonical” hydrogen bonds (i.e. those involving O-H and N-H donors) can, at least in some instances, promote folding when polar side chains are present at helix-helix interfaces^61–64^. This is because the loss of hydrogen bonding when the helices are separated is not compensated by the interactions between the polar side chains with the apolar lipid environment, which are not as favorable. The same argument should apply to the weak Cα−H hydrogen bonds, the signature trait of GAS _right_ dimers, but we still lack sufficient experimental evidence. Part of the issue is the typical strategy for measuring the contribution of hydrogen bonding in protein folding and interaction, which is based on mutating the polar side chains involved (such as Ser to Ala or Tyr to Phe, for example). This type of mutational strategy is unfeasible in the case of Cα–H∙∙∙O=C hydrogen bonds, because both donor and acceptor groups are part of the backbone. Therefore, measuring their contribution of Cα−H hydrogen bonds association has been exceedingly difficult^65,66^.

In this study, we advance these debates as they apply to the GAS_right_ motif, with a thermodynamic analysis of association of eight representative candidates from the original pool of 26 constructs we previously studied with TOXCAT^16^. Of these constructs, one formed higher oligomers whereas a second stood out as a clear outlier in the apparent relationship between ΔG° of association and TOXCAT signal. For the remaining six constructs, we found that there is a striking correspondence between their relative stability (ΔΔG°) in detergent and their predicted stability in the biological membrane of *E. coli* calculated from TOXCAT (Fig. 5c). Additionally, we found that when the free energy of association of the entire original pool of 26 constructs is back-calculated from TOXCAT using this apparent relationship, a significant correlation is observed with the ΔE calculated with CATM based on their predicted structural models (Fig. 6b). Therefore, the present data provide quantitative thermodynamic validation that is consistent with our previous findings^16^ and supports them.

The GAS_right_ motif is a versatile dimerization motif found both in structural proteins that require maximal stability, as well as in dynamic proteins, such as receptors, in which stability needs to be finely modulated to support their function. Combined, the present and previous studies provide insight into how sequence, structure, and energetic factors modulate this stability. The analysis of the predicted structural models identified significant trends that suggested that geometry can affect stability^16^. Specifically, shorter interhelical distances and crossing angles near −40° tend to produce the most stable dimers. In turn, these ideal geometries are favored by the presence of GxxxG motifs (as opposed to other GxxxG-like motifs, such as AxxxG or GxxxA). In particular, dimerization appears to become most effective when a Gly residue is present at the interfacial position designated as N1, forming a GxxxG with the Gly at C1 (Fig. 2*b*). None of these rules are stringent; it is possible for dimers containing GxxxG motifs to assume near-ideal geometry and, yet, their stability could be detuned if the shape of the residues at the dimer interface is not conducive to good packing ^28^. However, the analysis suggests that a geometric “sweet spot” exists, where weak hydrogen bonding and packing can be optimized, and which can be exploited by nature when maximal stability is required for function.

## Experimental Procedures

### Plasmid cloning

A gblock containing the soluble domain of Staphyloccocal nuclase (SN) (nuclease A-H124L from *Staphylococcus aureus*, amino acids ALA1-GLN149) fused to the TM domain of GpA (amino acids: E72-R96) through a flexible linker (amino acids: TSGG[SGGG]_2_SGGS) was inserted into a pet28a plasmid by restriction free cloning^67^. The construct included a C-terminal His-tag for purification. Mutations to the soluble domain SN to obtain S3C, A60C, and A112C were introduced using either QuickChange mutagenesis or double primer mutant method^68^. Restriction free cloning^67^ was used to replace the TM domain of GpA with the TM domain of the 8 GAS_right_ constructs obtained from the TOXCAT plasmid. All protein sequences are reported in supplementary Table S3. The following plasmids have been deposited on the AddGene repository with their respective accession numbers: glycophorin A construct pSN-A112C-GpA: #TDB; glycophorin A, G83I mutant construct, pSN-A112C-GpA-G83I: #TBD; CP8B1 construct, pSN-A112C-CP8B1: #TBD; 1A32 construct, pSN-A112C-1A32: #TBD;. 1A31 construct, pSN-A112C-1A31: #TBD; MCL1 construct, pSN-A112C-MCL1: #TBD.

### Protein expression

Plasmids were transformed into BL21 (DE3) non-tunner cells and plated in LB agar plates containing 50 μg/ml of kanamycin and incubated at 37 °C overnight. Colonies were inoculated in ~3 mL of LB media containing 50 μg/ml of kanamycin and incubated in a shaker overnight at 37 °C. Overnight cultures were inoculated into ZYP-5052 autoinduction media^69^ (1 ml of overnight culture per 500 ml of autoinduction media) containing 400 μg/ml of kanamycin and incubated in a shaker at 37 °C until the culture reached approximately 0.8 OD_600_, at which point the temperature was lowered to 25 °C and cells were left to grow overnight. Cells were collected by centrifugation at 5,000 ×g for 10 min. Pellets were washed with cell wash buffer (50mM TrisHCL pH 7.9, 100mM NaCl), then centrifuged again at 5,000 ×g for 10 min. The pellets were flash frozen in liquid nitrogen and stored at −80 °C.

### Purification of constructs

Cell pellets were mixed with lysis buffer (50 mM Tris HCl pH 7.9, 5 mM EDTA, 1 mM PMSF, 1 mg/ml Lysozyme and 5 mM BME) at a ratio of 10 ml of buffer per gram of cells, then lysed via sonication. The lysate was centrifuged at 10,000 ×g for 15 min. The membrane fraction was then isolated via ultracentrifugation of the supernatant at ~185,500 ×g for 30 min. The membrane pellet was then resuspended with approximately 5 ml/g of cells in solubilization buffer (50 mM Tris HCl pH 7.9, 1 M NaCl, 18 mM *n*-decyl-β-D-maltopyranoside and 10 mM TCEP), and left rotating overnight at 4 °C.

The membrane fraction was centrifuged at 5,000 ×g for 10 min. The supernatant was mixed with approximately 1 ml of Ni-NTA per gram of cells and equilibrated in solubilization buffer (50mM TrisHCl pH 7.9, 200mM NaCl, 5.4mM DM and 1mM TCEP). The sample was bound to the resin by rotating at 4 °C for at least 2 hours. The resin was loaded onto a gravity column and washed with wash buffer (50 mM Tris HCl pH 7.0, 200 mM NaCl, 5.4 mM DM, 1mM TCEP and 30 mM Imidazole). The purified protein was eluded with elution buffer (50 mM Tris HCl pH 7.0, 200 mM NaCl, 5.4 mM DM, 1mM TCEP and 200 mM imidazole). Samples were loaded on a 10DG desalting gravity column to remove the imidazole. The desalted fractions containing protein were then ultracentrifuged at 100,000 ×g for 30 min at 4 °C to remove possible aggregates. The final protein concentration was determined via UV-Vis spectroscopy using calculated extinction coefficients (supplementary Table S3).

### Labeling of chimeras with Cy3-maleimide or Cy5-maleimide

The constructs were labeled on-column with either Cy3-maleimide or Cy5-maleimide. First, the protein samples were mixed with Ni-NTA equilibrated with buffer (50mM TrisHCl, pH 7.0, 200 mM NaCl, 5.4mM DM and 1mM TCEP) and bound in batch at 4 °C for at least 2 hours. The resin was loaded on a gravity flow column and the flow through was collected. The resin was then washed with nonreducing wash buffer (50 mM TrisHCl pH 7.0, 150 mM NaCl, and 5.4 mM DM). The resin was then mixed with labeling buffer (50 mM Tris HCl pH 7.0, 150 mM NaCl, 5.4 mM DM and 0.1 mM fluorophore) using a 15-fold excess of fluorophore to protein, and incubated for 15 min. Samples were quenched for 5 min by mixing 100 mM L-cysteine in a 100-fold excess with respect to fluorophore. The column was washed with 75 ml of labeling wash buffer (50 mM Tris HCl pH 7.0, 150 mM NaCl, 5.4 mM DM and 30 mM Imidazole) and the constructs were eluded using labeling elution buffer (50 mM Tris HCl pH 7.0, 150 mM NaCl, 5.4 mM DM and 200 mM Imidazole). Purity was assessed by SDS-PAGE.

The most concentrated fractions were collected and dialyzed twice in 200 ml of dialysis buffer (50 mM Tris HCl, pH 7.0, 150 mM NaCl, and 5.4 mM DM) using a dialysis bag with a molecular weight cutoff of 12-14 kD. The samples were then ultracentrifuged at 100,000 ×g for 30 min at 4 °C to get rid of possible aggregates. The absorption spectra of the samples was measured to calculate the protein concentration *P* using the following equation:

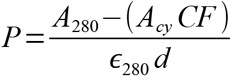

where A_280_ represents the absorbance at 280 nm, *A*_cy_ correspond to the absorbance at 553 nm for Cy3-maleimide or at 653 nm for Cy5-maleimide, *є*_280_ is the molar absorptivity for the corresponding construct, *d* is the cuvette path length (1 cm), and *CF* is the correction factor to account for fluorophore absorbance at 280 nm (0.04 for Cy5 and 0.1 for Cy3). To calculate the labeling efficiency *E_cy_* the following equation was used:

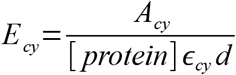

where *є*_cy_ represents the molar absorptivity for Cy3 (150,000 M^−1^cm^−1^) or Cy5 (250,000 M^−1^cm^− 1^). The labeling efficiencies for all constructs are reported in supplementary Table S4.

### Sample preparation for FRET experiments in detergent

Samples at different mole fractions (moles of protein/moles of detergent), were prepared by mixing 1.3 μM of Cy5-maleimide labeled protein with 1.3 μM of Cy3-maleimide labeled protein for a constant protein concentration of 2.6 μM in a buffer containing 50 mM TrisHCl pH 7.0, 150 mM NaCl, and varying concentrations of DM detergent such that mole fraction concentrations from ~1×10^−3^ to ~1×10^−5^ were covered. The mole fractions were calculated using total protein concentration, irrespective of the labeling efficiencies of the samples. In addition equivalent donor-only and acceptor-only samples were produced, each with a protein concentration of 1.3 μM. Samples were equilibrated at 4 °C in dark tubes. Fluorescence emission spectra were collected at various time points to check sample equilibration.

### FRET experiments

Emission spectra measurements were collected in a Tecan infinite M1000 pro plate reader. Samples were excited at 500 nm and emission spectra were collected from 505 nm to 800 nm every 1 nm step. The acceptor-only sample emission spectra were subtracted from the FRET samples emission spectra. Then the data was normalized by dividing every data point by the isosbestic point (640 nm) (supplementary Fig. S1). Equilibrium constants of dissociation were obtained by fitting the normalized fluorescence intensity at 670 nm (*NF*_670_) as a function of the protein:detergent mole fraction (*χ_T_*) using equation 1 (derived in supplementary material):

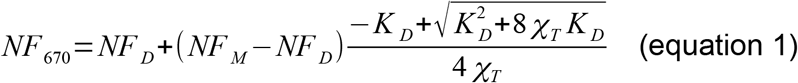

Three parameters were derived from the fitting: *NF_D_*, the normalized fluorescence intensity at the limit in which the samples are fully dimeric (i.e., at infinite concentration), *NF_M_*, the normalized fluorescence baseline at the limit in which the samples are fully monomeric (at infinite dilution), and *K_D_*, the desired dissociation equilibrium constant. The *NF_M_* and *NF_D_* parameters were fit as global variables, whereas each *K_D_*, was fit individually to their respective construct’s data.

The fraction of dimer *ϕ_D_* as a function of protein:detergent mole fraction *χ_T_* and dissociation constant *K_D_* was calculated with the following equation:

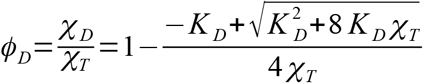

The dissociation free energy was determined from *K_D_* using the following equation:

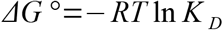

where *R* is the gas constant, *T* is the temperature. Statistical analysis was performed using the R software^70^, with the aid of the tidyverse^71^ and drc^72^ packages.

### Determination of the oligomeric state

Stoichiometry experiments were performed as previously described^35^ to confirm the oligomeric state of the constructs. Briefly Cy3-maleimide and Cy5-maleimide labeled protein samples were mixed at different donor:acceptor ratios from 10:90 to 90:10 with 10% increments while keeping the total protein concentration constant. The same was done for Cy3-maleimide and unlabeled protein samples. All samples were prepared at a mole fraction of 4.8×10^−4^. The relative donor quenching *Q* was obtained using the following equation:

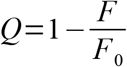

where *F* is the experimental quenched fluorescence for a specific donor:acceptor ratio, and *F*_0_ is the experimental unquenched fluorescence for the same amount of donor. To estimate the oligomeric state, the relative donor quenching *Q* was plotted as a function of donor fraction *P_D_*, and fit to the following equation:

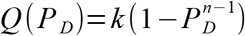

to obtain the oligomeric state *n*. The parameter *k* is defined by the following equation:

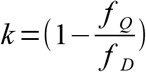

where *f_Q_* represents the molar fluorescence of the quenched donor, and *f_D_* represents the molar fluorescence of the donor in the absence of acceptor. The data was compared with theoretical model corresponding to different integer values of *n* (2, 3 and 4) and the oligomeric state was evaluated by examining the model with best correspondence.

## Abbreviations

TM: transmembrane
GpA: glycophorin A
FRET: Förster resonance energy transfer
SN: staphylococcal nuclease
TMD: transmembrane domain
DM: *n*-decyl-β-D-maltopyranoside
OD_600_: optical density at 600 nm
VDW: van der Waals
WT: wild type

## Acknowledgments

This work was supported by National Institutes of Health (NIH) grant R35-GM130339 to A.S. and National Science Foundation grants CHE-1710182 to A.S. and DMS-1661900 to Q.C.. G.D.V. acknowledge the support of a SciMed GRS Fellowship.

## SUPPLEMENTARY INFORMATION

**Table S1.** Initial labeling efficiencies of the constructs used for the screening of the labeling position on Staphyloccocal nuclase (SN).

| Construct | Cy5-maleimide (%) | Cy3-maleimide (%) |
| --- | --- | --- |
| SN-A112C-GpA-TM | 79.8 | 77.1 |
| SN-A112C-GpA-G83I-TM | 72.4 | 70.1 |
| SN-A60C-GpA-TM | 65.4 | 76.4 |

**Table S2.** The found relationship between TOXCAT and fraction of dimer was used to predict the standard free energy of the 26 constructs that we previously studied using TOXCAT and CATM (Predicted ΔG°). The relative free energy in the E. coli inner membrane was obtained with respect to poly-Leu GpA (ΔΔG°_poly-Leu GpA_).

| TM | Predicted $\Delta G^\circ$ | $\Delta\Delta G^\circ_{\text{poly-Leu GpA}}$ |
| --- | --- | --- |
| ROMO1 | -4.39 | 0.31 |
| K132L | -4.24 | 0.15 |
| CAD12 | -4.15 | 0.06 |
| GLPA | -4.08 | 0 |
| SEM6B | -4.06 | -0.02 |
| STAB1 | -4.03 | -0.05 |
| TNR12 | -4.03 | -0.05 |
| F159B | -3.96 | -0.13 |
| CP8B1 | -3.77 | -0.32 |
| 1A32 | -3.75 | -0.33 |
| OPA1 | -3.64 | -0.45 |
| ITA2 | -3.54 | -0.54 |
| SHSA7 | -3.39 | -0.69 |
| EPCR | -3.27 | -0.82 |
| FCRL4 | -3.27 | -0.82 |
| MGT5B | -3.21 | -0.88 |
| 1A31 | -3.08 | -1.01 |
| TNR1B | -3.05 | -1.03 |
| STBD1 | -3.05 | -1.03 |
| MUC3A | -2.71 | -1.37 |
| 1A32-2 | -2.67 | -1.41 |
| COX14-2 | -2.67 | -1.41 |
| 1A31-2 | -2.53 | -1.56 |
| TNR1B2 | -2.47 | -1.61 |
| MCL1 | -2.47 | -1.61 |
| ANPRC | -2.41 | -1.67 |

**Table S3.**
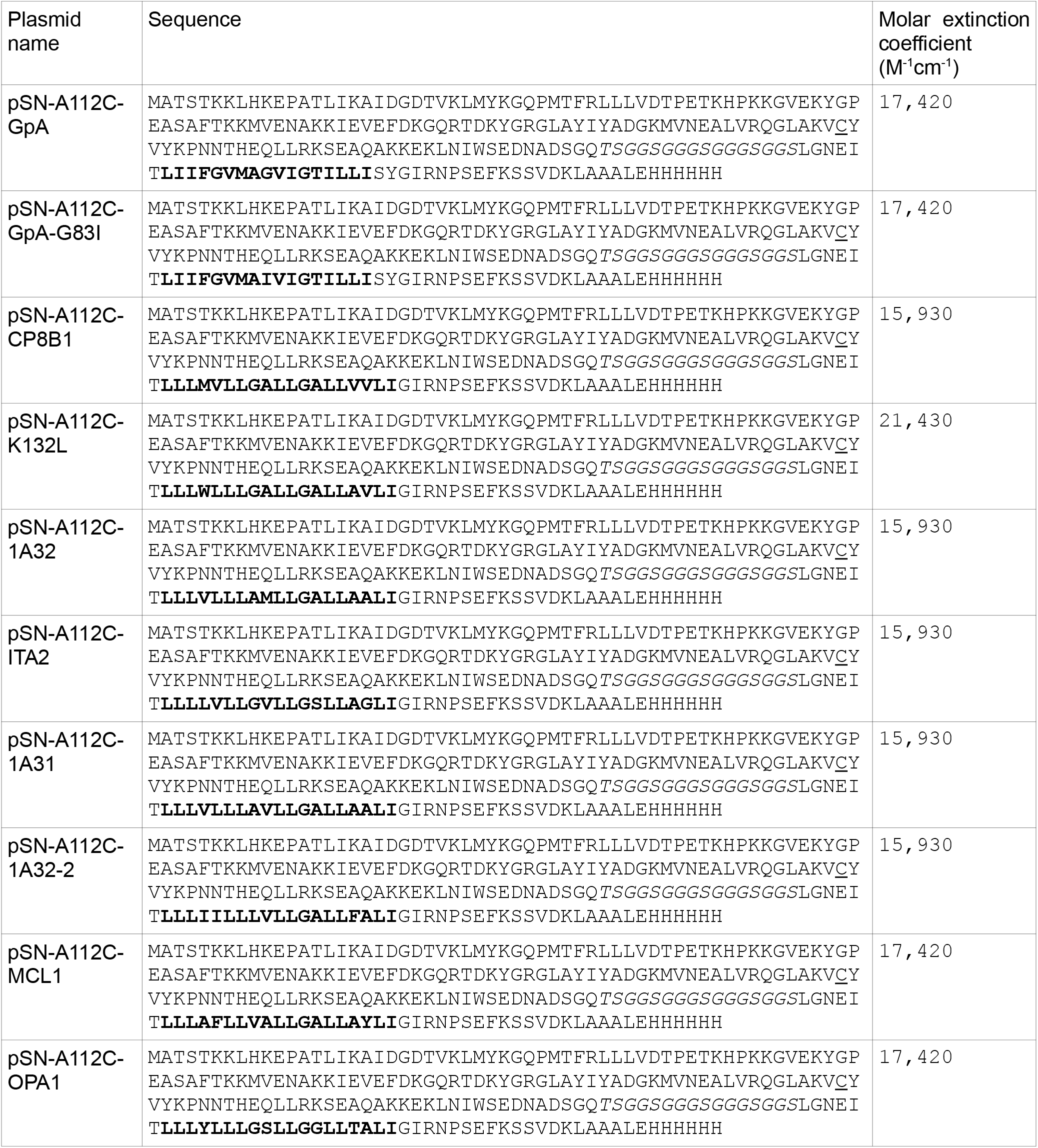
The amino acid sequence of each of the constructs used is presented, with the transmembrane region in bold. The flexible linker is in italics. The single Cys position introduced for fluorophore labeling (A112C) is underlined. The extinction coefficient used to determine the protein concentration for each of the constructs is also presented.

**Table S4.** Calculated labeling efficiencies for all different purified constructs.

| Construct | Cy5 (%) | Cy3 (%) |
| --- | --- | --- |
| SN-GpA | 68 | 87 |
| SN-G83I | 77 | 80 |
| SN-ITA2 | 74 | 80 |
| SN-ITA2 | 90 | 95 |
| SN-K132L | 76 | 77 |
| SN-K132L | 74 | 77 |
| SN-OPA1 | 71 | 85 |
| SN-OPA1 | 71 | 83 |
| SN-1A32-2 | 93 | 95 |
| SN-1A32 | 90 | 97 |
| SN-MCL1 | 60 | 68 |
| SN-CP8B1 | 72 | 78 |
| SN-1A31 | 72 | 85 |

### Derivation of the equation used for global fitting of the fluorescence data to obtain the dissociation constants

To derive the equation to fit the normalized fluorescence intensity at 670 nm (*NF_670_*) as a function of mole fraction to determine the equilibrium dissociation constant *K_D_*, we started from the following relationships:

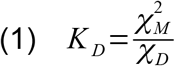

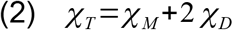

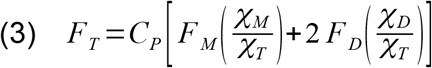

Where *χ_M_* is the monomer mole fraction, *χ_D_* is the dimer mole fraction, *χ_T_* is the total protein mole fraction, *F_T_* is the total fluorescence, *F_D_* is the fluorescence intensity of a fully dimeric sample, *F_M_* is the fluorescence intensity of a fully monomeric sample and *C_P_* is the total protein concentration use in the sample.

Rearranging (1) for *χ_D_* we get:

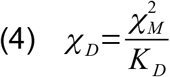

Using (4), we eliminate *χ_D_* from (2):

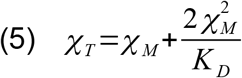

Rearranging (5) we get:

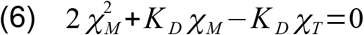

Solving the quadratic equation (6) for *χ_M_* we get:

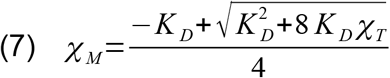

Rearranging (2) for *χ_D_* we get:

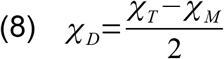

Using (8), we eliminate *χ_D_* from (3):

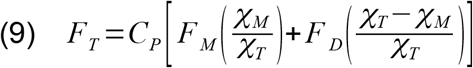

Rearranging (9) we get:

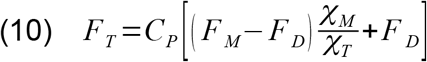

Using (7), we eliminate *χ_M_* from (10):

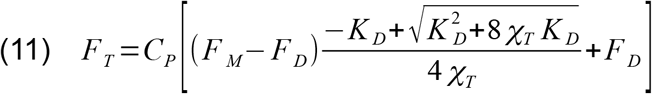

At the isosbestic point (640 nm, in our case) the fluorescence of the donor and acceptor samples are equal by definition, therefore:

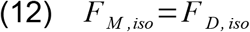

and thus, at this wavelength, the quadratic term drops and the total fluorescence in (11) simplifies to:

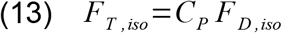

Dividing the fluorescence of the wavelength of interest (11) by the fluorescence at the isosbestic point (13) cancels out the protein concentration *C_P_*, resulting in the equation:

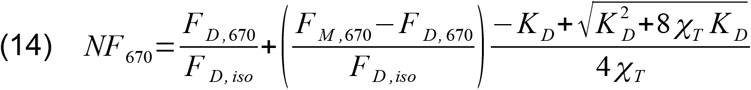

where *NF*_670_ refers to the normalized fluorescence intensity at the wavelength of interest (670 nm, in our case).

If we now redefine the normalized fluorescence of the fully monomeric (*NF_M_*) and dimeric samples (*NF_D_*) as the ratio of the fluorescence at 670 nm and the fluorescence at the isosbestic point:

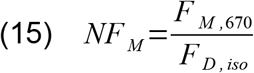

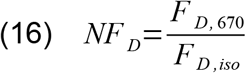

and substitute (15) and (16) in (14), we obtain the final form of the equation (equation 1):

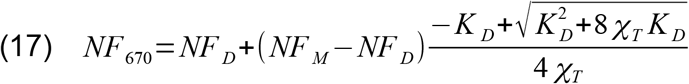

**Fig. S1.**
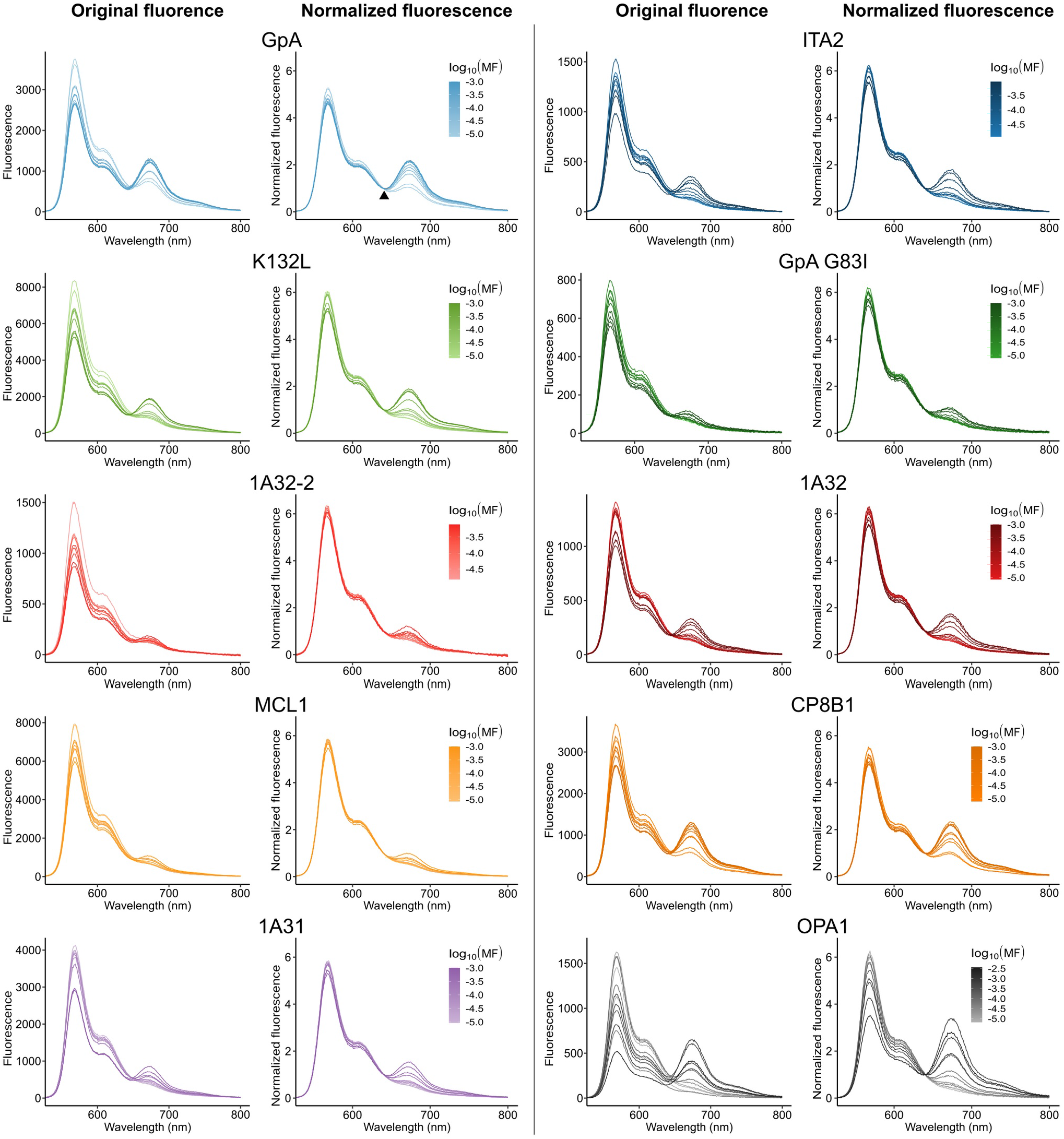
The original emission spectra of the different samples and their normalized version using the isosbestic point (640 nm for the FRET pair used, marked with the arrow in the GpA Normalized Fluorescence panel). The normalization was introduce to correct for small changes in signal that are unrelated to the transfer of energy between the donor and acceptor fluorophores, such as small errors in the estimate of concentration.

**Fig. S2.**
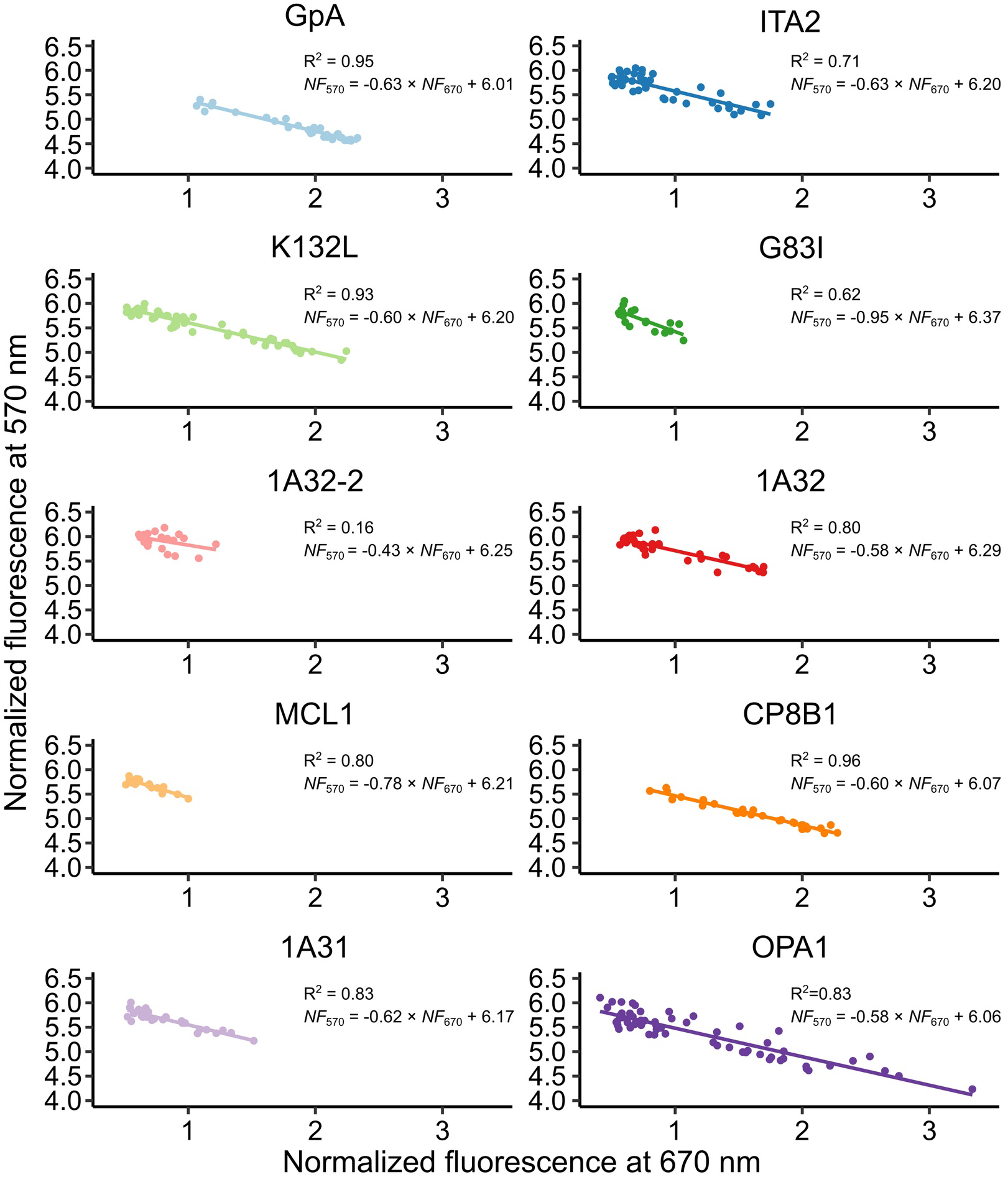
Donor quenching at 570 nm vs acceptor emission at 670 nm. The expected inverse linear relationship between donor quenching and acceptor sensitization is observed. For all constructs that cover the transition from mostly monomeric to mostly oligomeric (GpA, ITA2, K132L, 1A32, CP8B1, and OPA1), consistent slopes and intercept values are observed. The slopes corresponds to the average decrease in the normalized fluorescence of the donor per unit increase in acceptor normalized fluorescence. The intercepts are the theoretical emissions for a completely monomeric (zero FRET) sample. OPA1 displayed a larger emission range when compared to the other constructs suggesting it may be forming higher order oligomers. The low R^2^ for 1A32-2, and all the other weakly associating constructs (G83I, 1A31, MCL1) is expected. The data points for these nearly monomeric constructs are concentrated in a short range in the upper left corner, in the low-FRET region. As a result, the measurements do not contain sufficient spread for accurate linear regression.

**Fig. S3.**
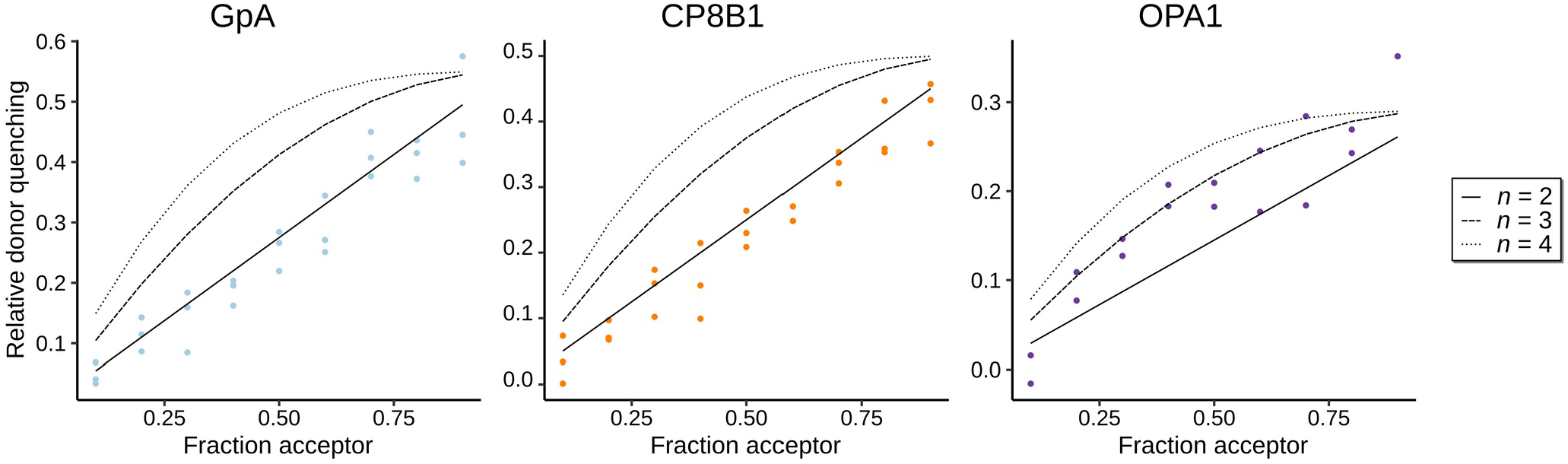
Stoichiometry experiments for the two outlier constructs CP8B1 and OPA1, and the control GpA. The experiments were carried out by varying the donor-labeled:acceptor-labeled construct ratio from 10:90 to 90:10 with 10% increments at a mole fraction of 4.8 × 10^−4^. Both GpA and CP8B1 have a good correspondence with the linear theoretical model (continuous line), indicating they associate as dimers (*n*=2). OPA1 corresponds better to an *n*=3 model (dashed line), indicating it is forming a higher-oligomer, likely a trimer.

**Fig. S4.**
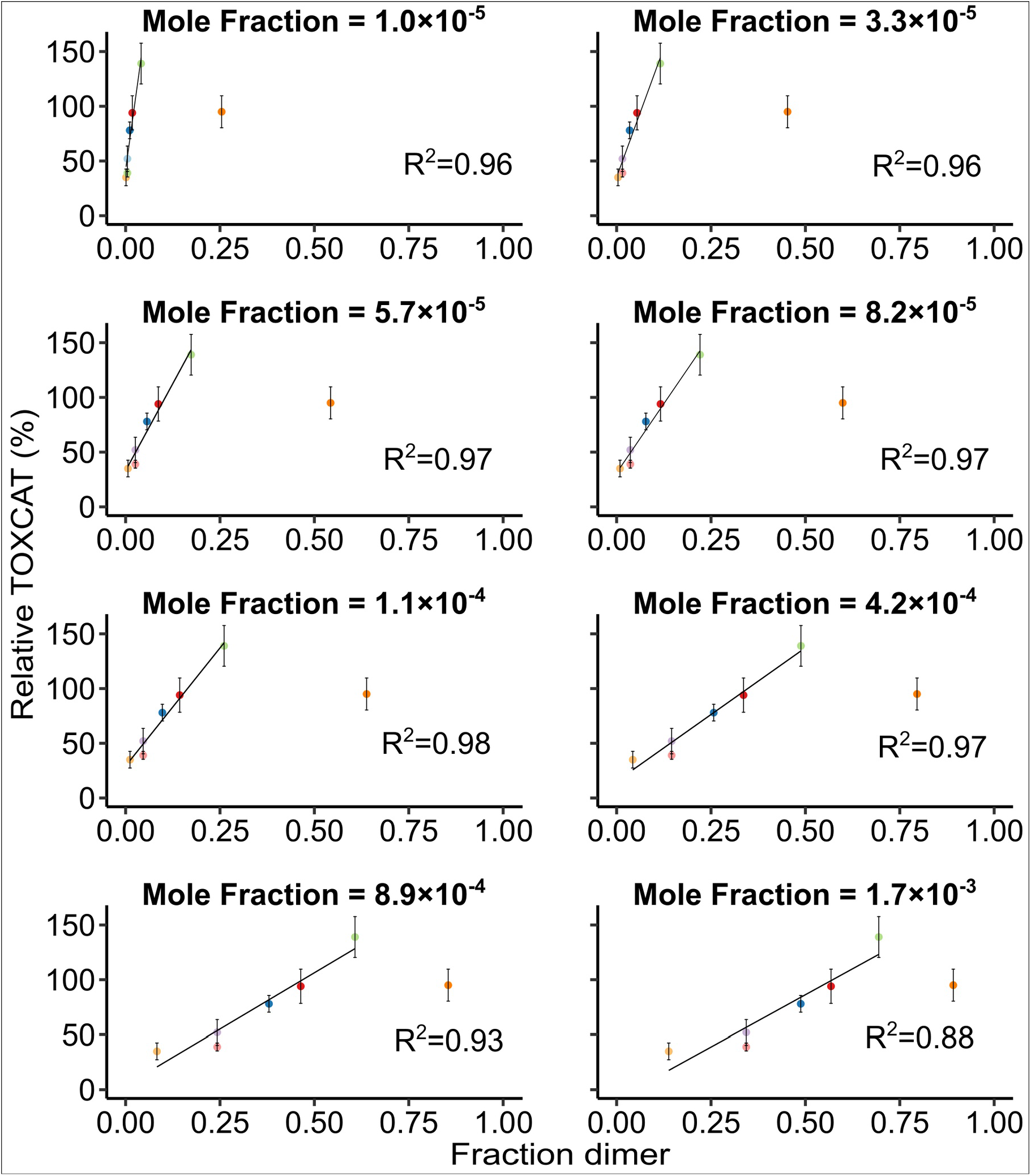
Comparison of the FRET results with the previous TOXCAT analysis, linear regression. Results of the linear regression analysis between TOXCAT and fraction of dimer calculated at differentprotein:detergent mole fractions to identify the concentration regime at which the linear relationship between dimer fraction in detergent and TOXCAT (taken as proxy for dimer fraction in the membrane) is maximized. The clear outlier CP8B1 was not included in this linear fit.

